# Riding the wave of genomics, to investigate aquatic coliphage diversity and activity

**DOI:** 10.1101/476168

**Authors:** Slawomir Michniewski, Tamsin Redgwell, Aurelija Grigonyte, Branko Rihtman, Maria Aguilo-Ferretjans, Joseph Christie-Oleza, Eleanor Jameson, David J. Scanlan, Andrew D. Millard

## Abstract

Bacteriophages infecting *Escherichia coli* have been used as a proxy for faecal matter and water quality from a variety of environments. However, the diversity of coliphages that are present in seawater remains largely unknown, with previous studies largely focusing on morphological diversity. Here, we isolated and characterised coliphages from three coastal locations in the UK and Poland. This revealed a surprising genetic diversity, with comparative genomics and phylogenetic analysis of phage isolates facilitating the identification of putative new species within the genera *RB69virus* and *T5virus* and a putative new genus within the subfamily *Tunavirinae*. Furthermore, by combining this genomic data with proteomic and host range analyses a number of phage structural proteins were identified, one of which is likely to be responsible for the observed differences in host range.

## Introduction

Bacteriophages are a key component of microbial communities playing important roles such as increasing the virulence and driving the evolution of their bacterial hosts, and influencing major biogeochemical cycles (see (1–3) for reviews). It is estimated that there are 10^31^ viruses in the biosphere with each millilitre of seawater containing millions of these viruses (1, 2) largely infecting the numerically dominant bacterial genera *Synechococcus*, *Prochlorococcus* and SAR11 (5–11). Culture- and metagenomics-based approaches have shed much light on their genetic diversity (12–16) including the description of several previously unknown phage groups that are widespread in the environment (9, 10, 17–19).

In the context of marine systems, bacteriophage infecting *Escherichia coli*, so-called coliphage, have perhaps received less attention even though they have been widely studied as a proxy for drinking water quality and the presence of faecal coliforms and enteric viruses (20–23). Thus, much is known about how the use of different *E. coli* strains or growth media used can lead to variable estimates of phage abundance (24–26) and this has resulted in global standards for using coliphages as a measure of water quality (27). For assessment of water quality these standards rely on the use of *E. coli* C strains derived from ATCC13706, which has been shown to detect increased titres over *E. coli* B and *E. coli* K12 derivatives (26). A criticism of the use of coliphages as indicators of water quality has been the reproduction of coliphages in the environment which will increase abundance estimates (28). Whilst the consensus seems to be that coliphage replication is not a significant issue (24), more recent research provides evidence that coliphages may well replicate in the environment (29).

Regarding the diversity of coliphages found in seawater, studies have largely focused on morphological diversity (29–32), assessing the number and range of *E. coli* hosts they can infect. This has shown that many coliphages have a broad host range, with detection of coliphages comprising members of the *Siphoviridae* and *Myoviridae* families off the Californian (29) and Brazilian coasts (31) but with *Siphoviridae* being the most frequently observed taxa (31).

Coliphages in general are one of the most sequenced phage types with ~450 complete phage genomes within Genbank, isolated from a variety of sources including animal faeces (33–36), human faeces (37), urine (38), clinical samples (39) river water (40), agricultural surface waters (41), lagoons (42), sewage (43) and animal slurries (34). However, as alluded to above, much less is known about the genetic diversity of coliphages in seawater. To begin to resolve this we isolated coliphages from three locations in the UK and Poland and undertook genomic and proteomic characterisation of the isolated phages, to provide insights into their phylogenetic position and functional potential.

## Results

For all samples tested the titre of coliphage detected was extremely low, generally <1 pfu ml^-1^ (Table 1). A total of 10 phage were isolated and purified from three different seawater samples and one phage from a freshwater urban pond. These phage were purified and their genomes sequenced to assess their genomic diversity (Table 1). Coliphage genomes were first compared against each other using MASH in an all-versus-all approach, which revealed three groups of phages based on similarity to each other: group1 vB_Eco_mar003J3 and vB_Eco_mar004NP2; group2: vB_Eco_mar005P1, vB_Eco_mar006P2, vB_Eco_mar007P3 vB_Eco_mar008P4 and vB_Eco_mar009P5; group3: vB_Eco_swan01, vB_Eco_mar001J1 and vB_Eco_mar002J2. Each phage was then compared against a database of complete phage genomes using MASH.

**Table 1.** Locations of water samples, titre of coliphages detected and phage isolates from each location.

| Water Source | Titre | Phage Isolates | Date of isolation |
| --- | --- | --- | --- |
| Oliva Stream Estuary, Jelitkowo,<br>Gdansk, Poland | 0.28 pfu ml<sup>-1</sup> | vB_Eco_mar001J1 | 30.01.2017 |
|  |  | vB_Eco_mar002J2 | 30.01.2017 |
|  |  | vB_Eco_mar003J3 | 30.01.2017 |
| Martwa Wisla Estuary, Nowy<br>Port, Gdansk, Poland | 0.11 pfu ml<sup>-1</sup> | vB_Eco_mar004NP2 | 30.01.2017 |
| Swanswell Pool, Coventry,<br>United Kingdom | 0.0125 pfu ml<sup>-1</sup> | vB_Eco_swan01 | 08.12.2016 |
| Great Yarmouth, United<br>Kingdom | ND | vB_Eco_mar005P1 | 08.12.2016 |
|  |  | vB_Eco_mar006P2 | 08.12.2016 |
|  |  | vB_Eco_mar007P3 | 08.12.2016 |
|  |  | vB_Eco_mar008P4 | 08.12.2016 |
|  |  | vB_Eco_mar009P5 | 08.12.2016 |

Phages vB_Eco_mar005P1, vB_Eco_mar006P2, vB_Eco_mar007P3, vB_Eco_mar008P4 and vB_Eco_mar009P5 had greatest similarity to phages APCEc01 (KR422352) and *E. coli* O157 typing phage 3 (KP869101), neither of which are currently classified by the ICTV. To further investigate the phylogeny of these phages, the gene encoding the major capsid protein (*g23*) was used to construct a phylogeny, as it is widely used as a phylogenetic marker including being used previously to classify phages within the *Tevenvirinae* (44). The *g23* sequence for the four newly isolated phages (vB_Eco_mar005P1, vB_Eco_mar006P2, vB_Eco_mar007P3, vB_Eco_mar008P4 and vB_Eco_mar009P5) were identical, therefore only one copy was included in the phylogenetic analysis. The analysis placed the new phage isolates within a clade that contains APCEc01, *E.coli* O157 typing phage 3, HX01, vB_EcoM_JS09 and RB69 (Figure S1). The latter three of these form part of the genus *Rb69virus,* suggesting the newly isolated phages are also part of this genus (Figure S1).

The genomes of phages from the genus *RB69virus* were further compared together with phage phiE142, which has an ANI of ~94% compared to the new isolates in this study. The ANI of all phages was calculated and compared in an all-v-all comparison, and the newly isolated phages had an ANI of >95% to HX01, JS09 and RB69 suggesting they are representatives of one of these species based on current standards (45). In fact, with the exception of phiE142, all phages had an ANI >95% with at least one other phage (Figure 1). To further elucidate the evolutionary history of these phages a core gene analysis was carried out. In the process of doing this, it became apparent phiE142 was ~50 kb smaller than the other phages within this group. Furthermore, it lacks essential genes that encode the major structural proteins and small and large subunit terminase. Therefore, it was excluded from further analysis as it is incomplete despite being described as complete (46).

**Figure 1.**
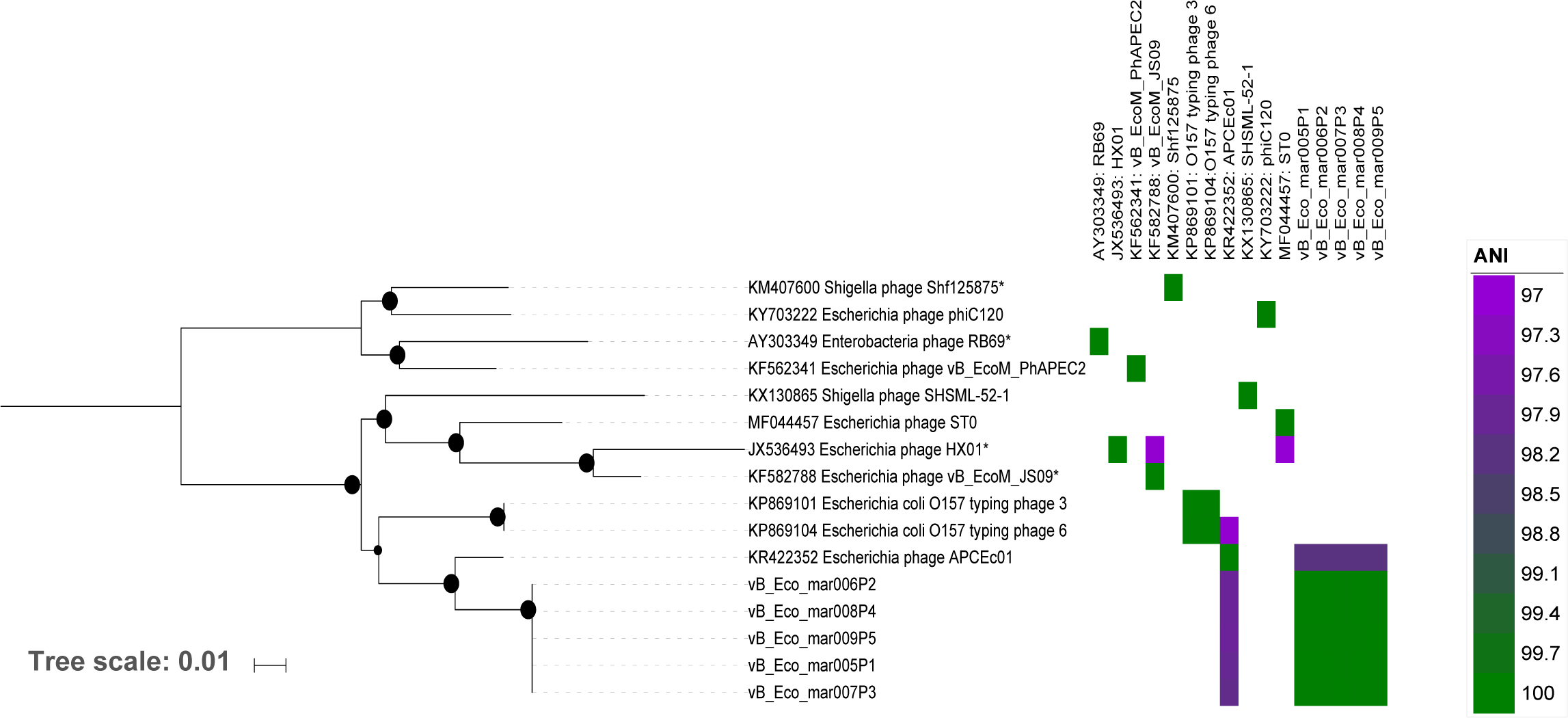
Phylogenetic analysis of phages within the genus *RB69virus.* The tree is based on the nucleotide sequence of nine concatenated genes using a GTR+F+ASC+R2 model of evolution, with 1000 bootstrap replicates using IQTREE (78). Current phage species as defined by the ICTV are marked with an *. Bootstrap values above 70% are marked with a filled circle, with the size proportional to the bootstrap value. The ANI value between phages is represented as a heatmap.

The core-genome of the genus *RB69virus* consisted of 170 genes, which accounted for 60.3-68.3 % of the total genes in each phage (Table S1). To further classify these phages, the GET_PHYLOMARKERS pipeline was used to identify suitable genes for phylogenetic analysis (47). Only 89 genes were identified that did not show signs of recombination when tested with Phi test (48). This test was carried out as recombination is known to result in inaccurate phylogenies and branch lengths (49). 86 of these passed further filtering to remove genes that were considered significant outliers using the KDETREES test (50). The resulting top nine genes (Table S1) as determined via GET_PHYLOGENIES were selected for phylogenetic analysis and a concatenated alignment was used for phylogenetic analysis (**Error! Reference source not found.**). Phylogenetic analysis placed the newly isolated phages in a clade with *Escherichia* phage APCEc01 (KR422352) further confirming they are they are part of the genus *RB69virus.*

Current taxonomy classifies RB69, HX01, JS09 and Shf125875 as four species within the genus *RB69virus* (Figure 1). This is based on the definition that phage species with ≥95% similarity based on BLASTn to another phage are the same species (45). The nucleotide identity between genomes was estimated using ANI by fragmentation of the genomes (51) rather than simple BLASTn comparison (45). Using an ANI value of >95% did not differentiate between phage species and maintained the current taxonomy, with each phage having an ANI >95% to multiple phages suggesting that *RB69virus* should contain only two species. Nevertheless, the phylogeny clearly supports multiple species within the *RB69virus* genus, suggesting a cut-off of 95% ANI may not be suitable (Figure 1). Consequently, if an ANI of >97% was used to differentiate species, this closely resembled the observed phylogeny (Figure 1). The higher ANI cut-off value discriminates between RB69 and Shf125875, which are currently classified as separate species. Furthermore, this will split the genus *RB69virus* into ten species, which are represented by Shf125875, phiC120, RB69, vB_EcoM_PhAPEC2, SHSML-52-1, STO, HX01, JS09, *E. coli* O157 typing phage 3 (strains *E.coli* O157 typing phage 6) and APCEc01 (including the five new isolates in this study). This suggests the five phages identified in this study are representatives of a new species within the genus *RB69virus*.

A similar approach was used for classification of the newly isolated phages vB_Eco_mar003J3 and vB_Eco_mar004NP2 which were most similar to phages within the genus *T5virus.* All phages that are currently listed as part of the genus *T5virus* were extracted from GenBank (April 2018). Initially, the gene encoding for DNA polymerase was used to construct a phylogeny, which has previously been used for the classification of phages within the genus *T5virus* (52) (Table S2). This confirmed that phages vB_Eco_mar003J3 and vB_Eco_mar004NP2 were related to other phages within the genus *T5virus* (Figure S2). Determination of the core-genome revealed 19 genes formed the core when using 90% identity for identification of orthologues using ROARY. However, when using this value and then applying the same filtering parameters as used for the genus *RB69virus*, no genes were deemed suitable for phylogenetic analysis. Therefore, an iterative process was used whereby the identity between proteins was lowered by 5% on each run of ROARY and the analysis repeated until a number of phylogenetic markers passed the filtering criteria, this was reached at a protein identity of 75%. At this point 44 core-genes were identified, of which only 14 passed further filtering steps (Table S2). The top nine markers as selected by the GET_PHYLOMARKERS pipeline were used for phylogenetic analysis (47).

Phylogenetic analysis on the selected marker genes confirmed that vB_Eco_mar004NP2 and vB_Eco_mar003J3 fall within the genus *T5virus* (Figure 2). Phage vB_Eco_mar004NP2 is a sister group to that of phage SPC35 (HQ406778) and vB_Eco_mar003J3 a sister group to that of phage LVR16A (MF681663) (Figure 2). Phage vB_Eco_mar004NP2 represents a new species within the genus *T5virus*, as it has <95% ANI with any other phage within the genus (45). For phage vB_Eco_mar003J3, it is not clear if the phage represents a new species. It has an ANI >95% with phages saus132, and paul149 which have recently been described as new species (52). However, these phages are not the closest group based on a phylogenetic analysis (Figure 2). When an ANI value of >97% is used then currently defined species are more congruent with the observed phylogenetic analysis, suggesting vB_Eco_mar003J3 is a novel species (Figure 2). Applying this threshold of 97% ANI across the entire genus would maintain the current species and create a total of 23 species across the genus.

**Figure 2.**
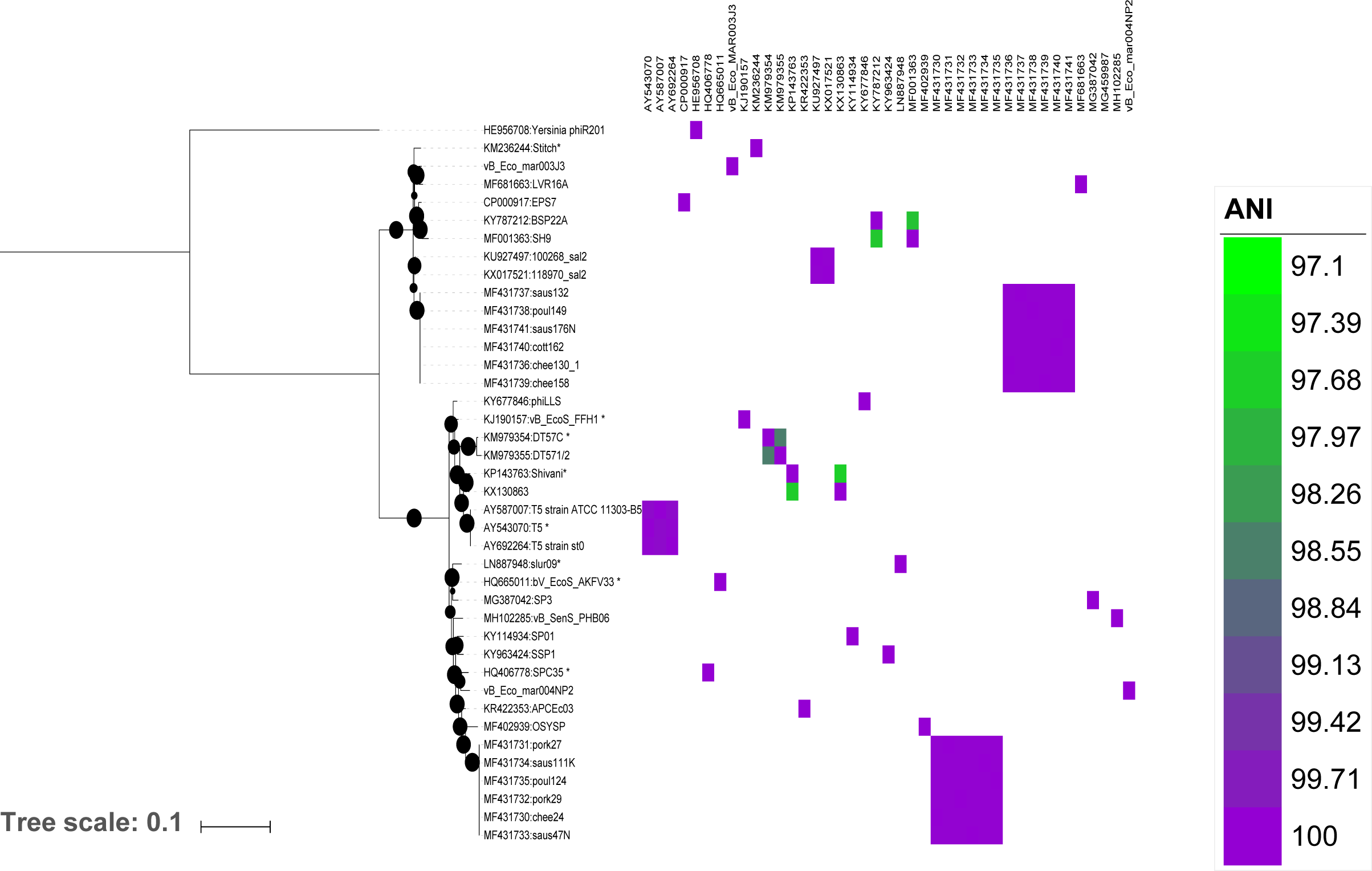
Phylogenetic analysis of phages within the genus *T5virus.* The tree is based on the nucleotide sequence of two concatenated genes using a GTR+F+ASC+R2 model of evolution, with 1000 bootstrap replicates using IQTREE (78). Current phage species as defined by the ICTV are marked with an *. Bootstrap values above 70% are marked with a filled circle, with the size proportional to the bootstrap value. The ANI value between phages is represented as a heatmap.

### Tunavirinae

Phages vB_Eco_mar001J1, vB_Eco_mar002J2 and vB_Eco_swan01 had greatest nucleotide sequence similarity to pSf-1 and SECphi27 which are members of the subfamily *Tunavirinae*. To classify the newly isolated phages, a phylogenetic analysis was carried out using the gene encoding the large subunit terminase that has previously been used to classify phages within the subfamily *Tunavirinae* by the ICTV (53). The analysis included all current members of the subfamily *Tunavirinae* (April 2018). The newly isolated phages vB_Eco_mar001J1, vB_Eco_mar002J2 and vB_Eco_swan01 form a clade with phages pSf-1, SECphi27 and Esp2949-1 (Figure S3). This clade is a sister to the clades that represent the previously defined genera of *KP36virus* and *TLSvirus*, thus clearly placing these new phages within the subfamily *Tunavirinae* (Figure S3).

To further clarify the phylogeny of these phages, again a core-gene analysis of all members of the subfamily *Tunavirinae* was carried out. Given these phage form part of a taxonomic sub-family, using ROARY with similarity cut-off values of 90% resulted, unsurprisingly, in the detection of no core genes. Therefore, an alternative method was used using an orthoMCL approach from within GET_HOMOLOGUES software (54). OrthoMCL based analysis identified a core of only nine genes, which were then filtered in the same manner as for the *RB69virus* and *T5virus* genera. A phylogeny was then constructed based on the concatenated alignment of four core-genes (Figure 3).

**Figure 3.**
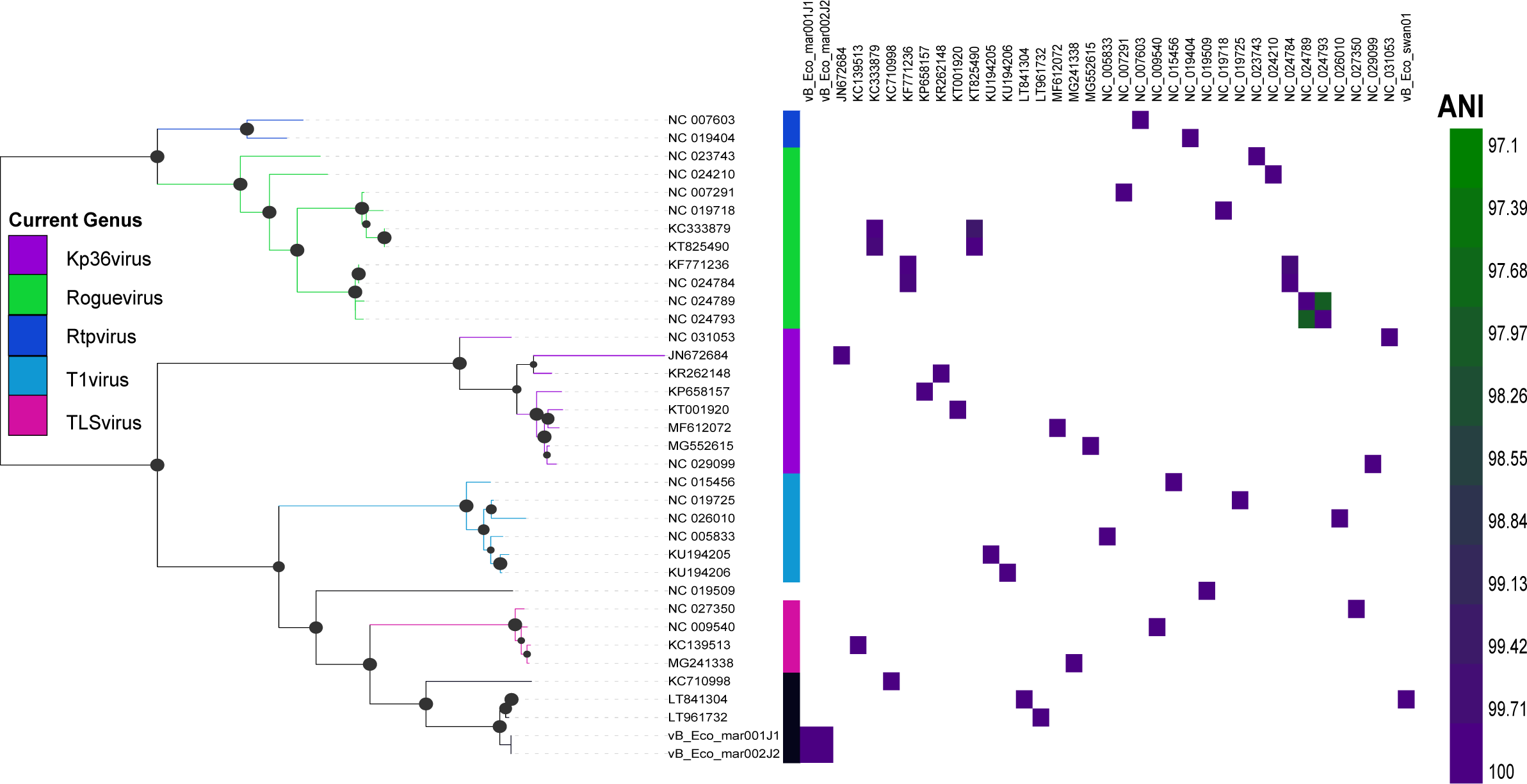
Phylogenetic analysis of phages within the subfamily *Tunanvirnae.* The tree is based on the nucleotide sequence of four concatenated genes using a GTR+F+ASC+G4 model of evolution, with 1000 bootstrap replicates using IQTREE (78). Current phage genera as defined by the ICTV are marked with the first coloured strip chart. Bootstrap values above 70% are marked with a filled circle, with the size proportional to the bootstrap value. The ANI value between phages is represented as a heatmap.

Phylogenetic analysis confirmed the previously defined genera within *Tunavirinae*, with the five genera of *Kp36virus*, *Roguevirus*, *Rtpvirus*, *T1virus* and *TLSvirus* also supported by good bootstrap support values (Figure 3). Furthermore, a clade which is sister to that of genus *TLSvirus* was identified with good bootstrap support comprising vB_Eco_mar001J1, vB_Eco_mar002J2, vB_Eco_swan01, SECphi27 (KC710998) and pSf-1 (NC_021331). Their clear separation from existing genera within the subfamily suggests this clade is a new genus. The phages within this putative genus all share an ANI >75% with other phages in the genus, compared to 60-70% ANI with phages in the other described genera within the *Tunavirinae*. All phages within the putative genus have a conserved genome organisation and share thirty orthologues. We propose that this clade represents a new genus and should be named *pSF1virus* after pSF-1, the first representative isolate. Furthermore, we propose the unclassified phage Esp2949-1 (NC_019509) is the sole representative of a new genus, as it doesn’t currently fit within currently defined genera. Phylogenetic analysis indicates that phages of the genus *TL1virus*, *TLSvirus*, *psF1virus* all have a common ancestor, with Esp2949-1 ancestral to phages in the genus *TL1virus* and *psF1virus*. (Figure 3). Comparative genomic analysis also supports this, with Esp2949-1 having <70% ANI to phages of the genera *TL1virus* or *TLSvirus,* its closest relatives. Phages within the putative genus *psF1virus* were further analysed to determine the number of species. Using a cut-off of 95% or 97% ANI, the genus will contain three species vB_Eco_swan01 (SECphi27, vB_Eco_swan01), vB_Eco_mar002J2 (vB_Eco_mar001J1, vB_Eco_mar002J2) and the orphan species pSF-1.

Phylogenetic analysis demonstrated that of the ten phages isolated, five represented novel species. A representative of each of these newly identified groups was further characterised both morphologically and physiologically. The representative phages were vB_Eco_swan01 and vB_Eco_mar002J2 (new species within the *Tunavirinae*), vB_Eco_mar003J3 and vB_Eco_mar004NP2 (new species within *T5virus*), and vB_Eco_mar005P1 (new species within *RB69virus*).

### TEM

TEM analysis confirmed they were all members of the order *Caudovirales* (Figure 4, Table 2), which contains all known tailed bacteriophages. Furthermore, phages vB_Eco_mar002J2, vB_Eco_mar003J3, vB_Eco_mar004NP2 and vB_Eco_swan01 were observed to have long non-contractile tails with a polyhedral head which are signatures of the family *Siphoviridae*. The length:width ratio further classified the phages within subgroup B1 (55). Phage vB_Eco_mar005P1 was also observed to have a polyhedral head, but with a long contractile tail, with tail fibres clearly observable which allows classification within sub group A2 within the *Myoviridae* (55) (Figure S4, Table 2).

**Table 2.**
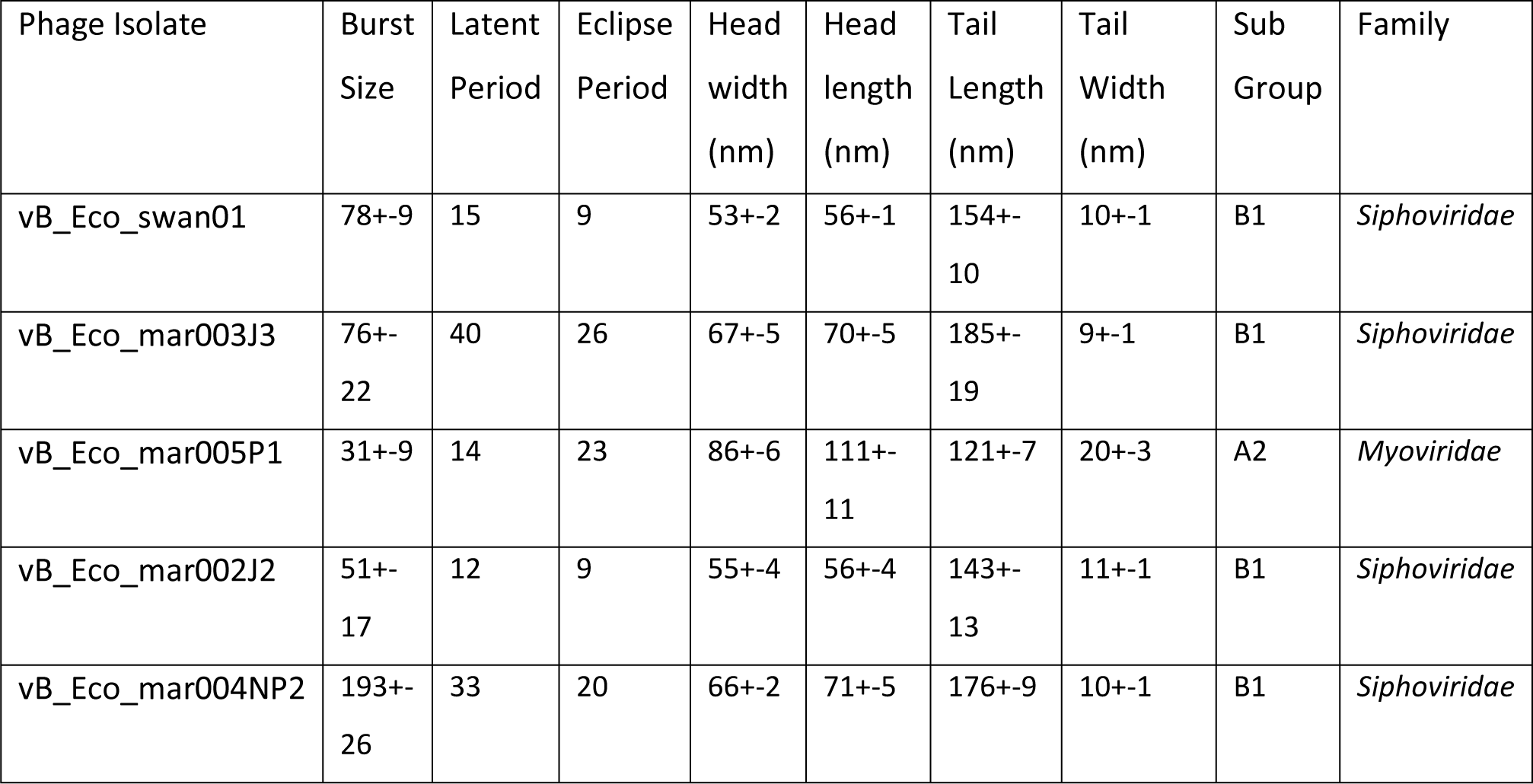
Morphological and lytic properties of representative phages.

| Phage Isolate | Burst Size | Latent Period | Eclipse Period | Head width (nm) | Head length (nm) | Tail Length (nm) | Tail Width (nm) | Sub Group | Family |
| --- | --- | --- | --- | --- | --- | --- | --- | --- | --- |
| vB_Eco_swan01 | 78+-9 | 15 | 9 | 53+-2 | 56+-1 | 154+-10 | 10+-1 | B1 | *Siphoviridae* |
| vB_Eco_mar003J3 | 76+-22 | 40 | 26 | 67+-5 | 70+-5 | 185+-19 | 9+-1 | B1 | *Siphoviridae* |
| vB_Eco_mar005P1 | 31+-9 | 14 | 23 | 86+-6 | 111+-11 | 121+-7 | 20+-3 | A2 | *Myoviridae* |
| vB_Eco_mar002J2 | 51+-17 | 12 | 9 | 55+-4 | 56+-4 | 143+-13 | 11+-1 | B1 | *Siphoviridae* |
| vB_Eco_mar004NP2 | 193+-26 | 33 | 20 | 66+-2 | 71+-5 | 176+-9 | 10+-1 | B1 | *Siphoviridae* |

**Figure 4.**
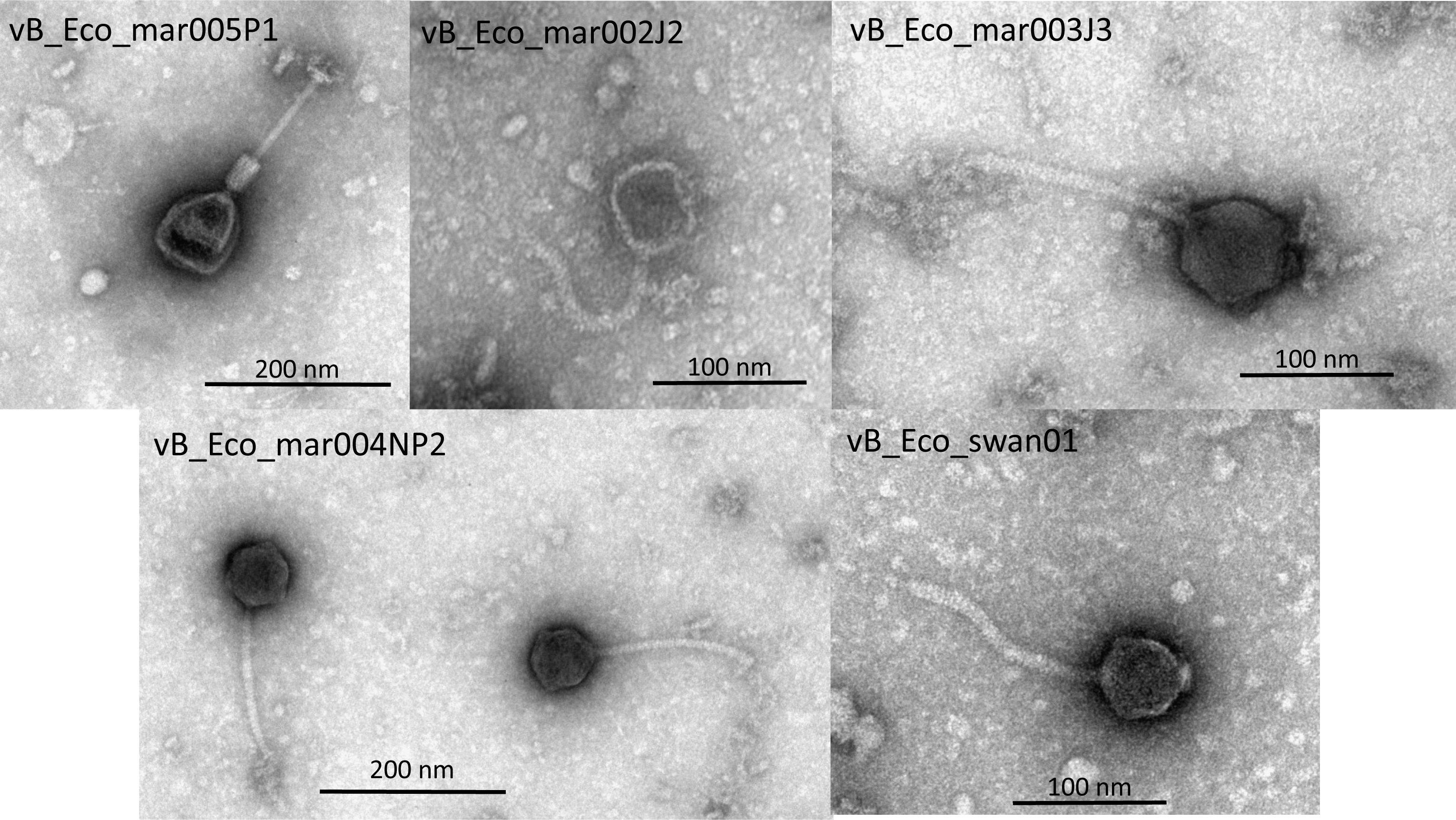
Morphology of phage isolates. Phages vB_Eco_swan01, vB_Eco_mar005P1, vB_Eco_mar002J2, vB_Eco_mar003J3, vB_Eco_mar004NP2 were stained with 2% (w/v) uranyl acetate and imaged in a JEOL JEM-1400 TEM with an accelerating voltage of 100 kV.

### Proteomic Characterisation

As with most phages the majority of the genes predicted within each genome encode for hypothetical proteins with unknown function. In order to identify further structural proteins or proteins that may be contained within the capsid, proteomic analysis of representative phages was carried out using electrospray ionization mass spectrometry (ESI-MS/MS). The number of identified proteins per phage was five, five, seven and eight for phages vB_Eco_mar005P1, vB_Eco_swan01, vB_Eco_mar003J3, and vB_Eco_mar004NP2 respectively (Table 3). This allowed the confirmation of two annotated structural proteins (SWAN_00017 and SWAN_00019) and the identification of a further three structural proteins (SWAN_00025, SWAN_00026, SWAN_00027). Based on the core-gene analysis this allowed annotation of orthologues of SWAN_00017, SWAN_00019, SWAN_00025 in vB_Eco_mar001J1, vB_Eco_mar002J2 and SECphi27, and SWAN_00026 and SWAN_00027 in vB_Eco_mar001J1 and vB_Eco_mar002J2.

**Table 3.**
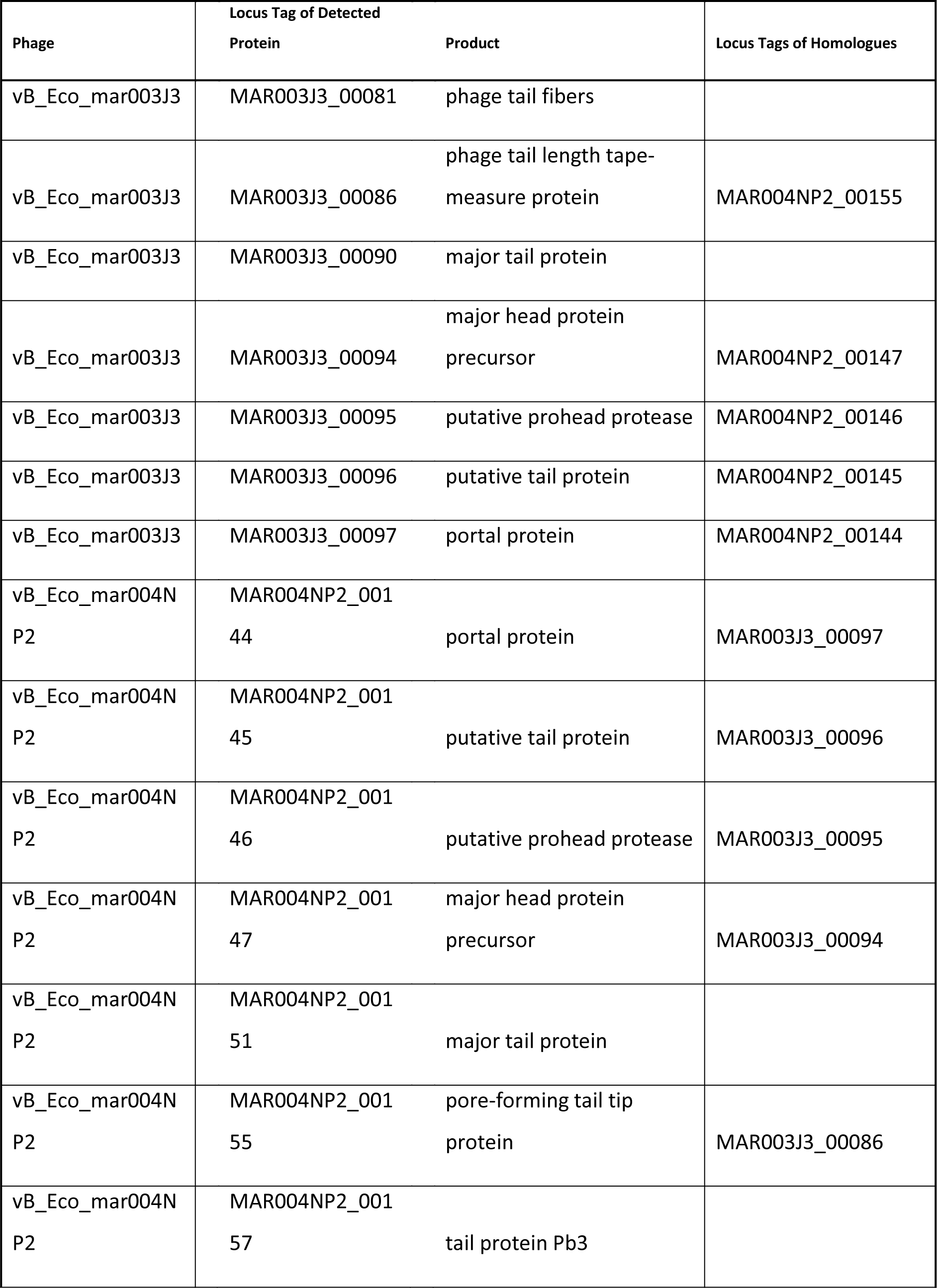
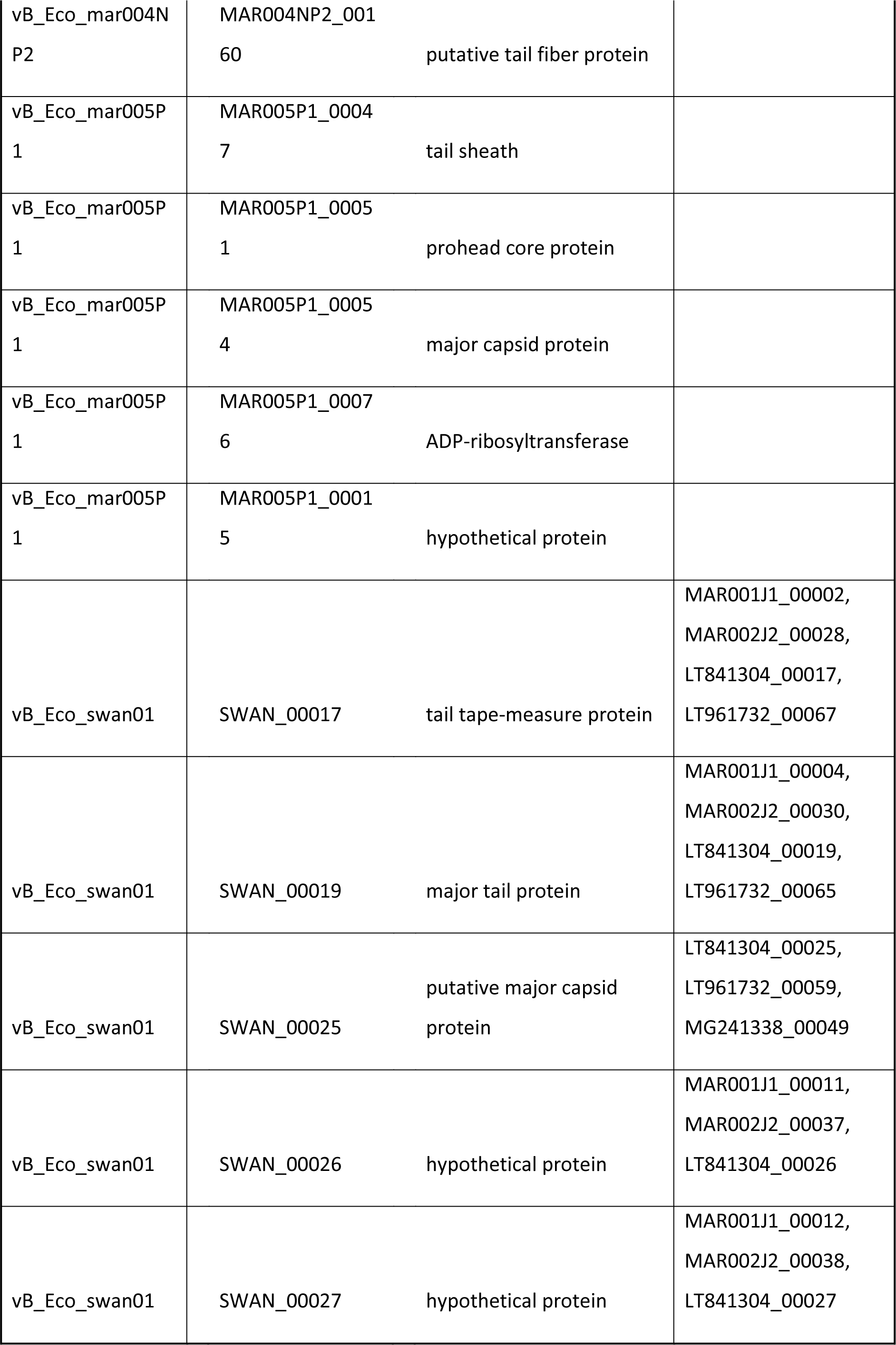
Proteomic analysis of phages vB_Eco_swan01, vB_Eco_mar005P1, vB_Eco_mar002J2, vB_Eco_mar003J3 and vB_Eco_mar004NP2.

For phage vB_Eco_mar005P1, five proteins were identified three of which confirmed annotations as structural proteins (MAR005P1_00047, MAR005P1_00051, MAR005P1_00054) all of which are core-genes to phages within the genus *RB69virus*, along with an ADP-ribosyltransferase protein (MAR005P1_00076) that is packaged within the phage capsid. An additional structural protein (MAR005P1_00015) was confirmed that was previously annotated as a hypothetical protein, which is also found in phages vB_Eco_mar005P1, vB_Eco_mar006P2, vB_Eco_mar007P3, vB_Eco_mar008P4 and vB_Eco_mar009P5.

Both phages vB_Eco_mar004NP2 and vB_Eco_mar003J3 are part of the genus *T5virus*, although distantly related. For phage vB_Eco_mar004NP2 eight proteins were detected that confirmed their annotation as various structural components of the capsid and tail (Table 3). For proteins MAR003J3_00086 and MAR003J3_00094-97 the orthologous proteins in vB_Eco_mar004NP2 were also detected. The proteins MAR004NP2NP2_00151, MAR004NP2_00157 and MAR004NP2_00160 were only detected in vB_Eco_mar004NP2. However, orthologous proteins were detected in vB_Eco_mar003J3 through core-gene analysis. The protein MAR003J3_00081 which is a putative tail fibre was only detected in vB_Eco_mar003J3, with no orthologue in vB_Eco_mar004NP2 based on core-gene analysis.

### Phage infection parameters

The burst size, latent period and eclipse period for representative phage isolates was also determined (Table 2). There was considerable variation in these parameters across all isolates, with burst size ranging from 31 (vB_Eco_mar005P1) to 192 (vB_Eco_mar004NP2) (Table 2). Similar variation was observed for the latent period varying from 12 min (vB_Eco_mar002J2) to 40 min (vB_Eco_mar003J3) whilst the eclipse period ranged from 9 min (vB_Eco_swan01 & vB_Eco_mar002J2) to 26 min (vB_Eco_mar003J3). For phages vB_Eco_mar003J3 and vB_Eco_mar004NP2 that are part of the same genus (T*5virus)*, there was considerable variation in all three parameters, with the burst size of vB_Eco_mar004NP2 (193) double that of vB_Eco_mar003J3 (76).

### Phage host range

The host range of representative phage isolates was determined using a range of bacterial hosts via a spot test assay (Table S4). Phylogenetic analysis highlighted that the isolated coliphages were often closely related to phages that are known to infect other Enterobacteriaceae, including *Klebsiella* and *Salmonella* (Figures 1, 2, 3). For this reason the host range of these phage was also tested against other Enterobacteriace. Phage vB_Eco_mar005P1 a representative of the genus *RB69virus* was only able to infect its host of isolation (*E. coli* MG1655), whereas phages of the genus *T5virus* and subfamily *Tunavirinae* were capable of infecting between five and eight strains (Table S4). Whilst vB_Eco_mar002J2 was found to infect the greatest number of strains (8), this was limited to strains of *E. coli*, *Klebsiella pneumoniae and Klebsiella oxytoca*, whereas vB_Eco_mar004NP2 could also infect *Salmonella typhimurium,* but fewer strains of *E. coli*.

### Detection in viral metagenomes

The presence of these new coliphage species in viral metagenomes was investigated using existing metagenomics databases. The Baltic virome dataset was chosen as it contains both DNA sequence data and RNA expression data (56). The abundance of representative phage species was determined by the stringent mapping of reads from the virome to representative genomes. *Synechococcus* phage Syn9 was also included, as it has previously been demonstrated to be present in this dataset (56). The overall coverage of each genome was low, with small numbers of reads mapping to each genome (Figure S4a). However, reads mapping to coliphage were found, although at far less abundance than cyanophages Syn9 (Figure S4a). We then searched for evidence of gene expression from these phages using transcriptomic datasets. The majority of samples showed expression of cyanophage Syn9 genes, as previously reported (56). In contrast, genes from coliphage NP2 and RB69 (Figure S4b) were only detected in samples GS852 and GS677, respectively. These samples, GS852 and GS677, were collected from low salinity surface waters (56). The reads mapping to coliphages were further analysed by BLASTn against the nr database. The only significant similarity in addition to the genomes they mapped against was to an un-annotated prophage region in five *E. coli* genomes, thus are likely transcripts from phages.

## Discussion

Using *E.coli* MG1655 we were able to isolate and characterise ten phages from coastal marine waters and one from a freshwater pond. The titre of coliphages in all water samples was extremely low (range 0.0125 pfu ml^-1^-0.28 pfu ml^-1^). This low abundance is lower than previous reports of coliphages that are present at an order of 1 x 10^2^ pfu ml in other coastal environments (57–59). This lower abundance may well be linked to water quality, as faecal contamination is known to be linked to coliphage abundance and/or the time of sampling. Only one sample point was collected, and previous work has found there are distinct seasonal patterns in coliphage abundance (59). Despite this low abundance, it was still possible to isolate coliphages to further characterise their genetic diversity, which was the focus of this study.

Given the small number of phages isolated and sequenced, there was a surprising amount of genomic diversity. Five species of coliphage were identified in the 10 phages isolated. The phages vB_Eco_mar005P1, vB_Eco_mar006P2, vB_Eco_mar008P4 and vB_Eco_mar009P5 were identical, with vB_Eco_mar007P3 only differing from the others by a single SNP. This similarity is probably due to the enrichment method, which has enriched for a single phage that has then proliferated in the enrichment and been re-isolated. Phages vB_Eco_mar001J1 and vB_Eco_mar002J2 also had identical genome sequences despite being independently isolated, and represent a novel species. The remaining phages vB_Eco_mar003J3, vB_Eco_mar004NP2, vB_Eco_swan01 were all unique and also represent new species.

Phages infecting *Escherichia* account for ~7% of all phages sequenced to date. To discover a novel genera from the sequencing of a small number of coliphages here further highlights the vast diversity of phages present in the environment and how much more is to be discovered. To accurately place phages in the context of current phage taxonomy, we identified core-genes and used the GET_PHYLOMARKERS pipeline to select the most appropriate gene for phylogeny reconstruction that do not show signs of recombination, and are thus likely to lead to inaccurate branch lengths (49). Our phylogenetic analysis of phage genomes using selected marker genes was congruent with current classifications of phage species. Some of these classifications are originally based on historical phenotypic data such as phage RB69 which cannot recombine with phage T4 and was classified as a separate species (60). Recently, this inability to recombine with phage T4 DNA was postulated to be caused by the arabinosyl modification of DNA in RB69, likely caused by a novel glucosyltransferase present in RB69 but not T4 (61). In this study, the gene thought to encode a putative arabinosyltransferase (61), was found to be core to all members of the genus *RB69virus*. Whether the phage isolated in this study also glycosylate their DNA in a similar manner to RB69 remains to be determined. However, the genes thought to be responsible for it are clearly a signature of this genus.

Whilst the phylogenetic analysis was congruent with currently defined species within the *T5virus* and *RB69virus* genera, combining this phylogenetic analysis with ANI data demonstrated that using an ANI value >95% was insufficient to delineate species that were congruent with the observed phylogeny when additional phage from this study, and those present in GenBank but having undefined species were added. Phages that form clearly distinct clades had an ANI >95% with phages outside of the phylogenetic clades. Thus, suggesting 95% ANI is insufficient to discriminate between species for some genera. We therefore suggest an ANI of 97% should be used to discriminate phage within the genera *T5virus* and *RB69virus*, which has previously been used for the demarcation of phage species within the genus *Seuratvirus* (62).

Proteomic analysis of the representative phages resulted in a relatively small number of proteins being detected per phage. Despite this, it was still possible to confirm the annotation of structural proteins and identify new structural proteins in phage vB_Eco_mar005P1 and vB_Eco_swan01. Combined with the core-gene analysis it confirmed the annotation of a large number of genes across all phage isolates as structural proteins. In addition, the detection of a ADP-ribosyltransferase in vB_Eco_mar005P1 suggests that the carriage of this protein is common to phages in the genus *RB69virus* and presumably acts similarly to the ADP-ribosyltransferase carried by phage T4, in modifying the host RNA polymerase for early gene transcription (63, 64). For phage vB_Eco_mar003J3 a putative tail fibre gene (MAR003J3_00081) was detected for which there is no orthologue in vB_Eco_mar004NP2.

The gene encoding MAR003J3_00081 is an orthologue of *ltfA* in phage DT57C and DT571/2 which with l*tfB* encode for L-shaped tail fibres that allow attachment to different O-antigen types. This arrangement of two genes encoding for the L-shaped tail fibres is different from T5 which encodes the L-shaped tail fibres in a single gene (65, 66). vB_Eco_mar003J3 contains orthologues of both *ltfA* and *ltfB*, suggesting that it too uses two gene products for L-shaped tail fibres, whereas vB_Eco_mar004NP2 only contains an orthologue of *ltfB* (MAR004NP2_00162) and does not contain an orthologue of the single gene used by T5 (*ltf*). Comparison of the genomic context of the region of *ltfB* in vB_Eco_mar004NP2 reveals two genes immediately upstream of *ltfB* that do not have orthologues in vB_Eco_mar003J3, one of which likely encodes a protein to form the L-shaped tail fibre with the product of *lftB*. Similarly, there are two genes upstream of *ltfAB* in vB_Eco_mar003J3 that are absent in vB_Eco_mar004NP2. However, immediately beyond this the genome contains 10 genes either side of these genes that are present in the same order in both genomes. Given the observed difference in host range between phages vB_Eco_mar003J3 and vB_Eco_mar004NP2, we speculate that it is the differences in this region that contains tail fibre genes that are likely responsible and contributes to the ability of vB_Eco_mar004NP2 to infect multiple genera of Enterobacteriaceae.

Differences in the properties of vB_Eco_mar003J3 and vB_Eco_mar004NP2 were also observed in terms of their replication parameters, with vB_Eco_mar004NP2 having a burst size (193) twice that of vB_Eco_mar003J3 (76). It has previously been reported that phage chee24 which is also part of the genus *T5virus*, has a burst size of 1000 and a latent period of 44 mins (52), whereas other phages of the *T5virus,* such as phage T5 and chee30 have burst size of ~77 and ~44 respectively, suggesting considerable variation within the genus.

In comparison, there was similar variation in the burst size of phages within the genus *RB69virus*, with vB_Eco_mar005P1 having a burst size that is very similar to the reported burst sizes of 31 for phage RB69, but smaller than the burst size of 96 for phage APCE01 (37). Whether the lytic properties of phages does correlate with phylogeny requires more data than is currently available and would require standardised growth conditions for like-for-like comparisons, as it is known differences in temperature can influence burst size.

Detection of reads from the Baltic virome using high stringency mapping suggests the coliphage isolated in this study can also be found in the Baltic Sea, albeit at low abundance. Given some of the samples used in the Baltic virome were collected from sources close to human habitation, detection of coliphages is not completely surprising. In contrast the detection of both coliphage and Syn9 transcripts in the meta-transcriptomics dataset was. Transcripts from a phage (Syn9) infecting a photosynthetic cyanobacteria would be expected in marine samples and has previously been reported for this Baltic virome (56). However, the detection of coliphage transcripts at two sites was surprising given coliphages are not thought to actively replicate in seawater (24).

## Conclusions

We have begun to elucidate for the first time the genomic diversity of coliphage within seawater, identifying phages that represent several novel taxa, further expanding the diversity of phages that are known to infect *E. coli*. Furthermore, the analysis and identification of core-genes and selection of genes suitable for phylogenetic analysis provides a framework for the future classification of phages in the genera *RB69virus*, *T5virus* and subfamily *Tunanvirinae*. We further suggest that an ANI of >95% is not suitable for the delineation of species within the genera *RB69virus* and *T5virus* and that a value of >97% ANI should be used. Characterisation of phage replication parameters and host range further reinforces that morphologically similar phage can have diverse replication strategies and host ranges. Whilst we are cautious about the detection of coliphage transcripts in seawater metatranscriptomes, the most parsimonious explanation is that coliphage are actively replicating, an observation that certainly warrants further investigation.

## Materials and Methods

*Escherichia coli* MG1655 was used as the host for both phage isolation and phage characterisation work. *E. coli* MG1655 was cultured in LB broth at 37°C with shaking (200 rpm). Seawater samples were collected from UK and Polish coastal waters (see Table 1), filtered through a 0.22 µm pore-size polycarbonate filter (Sarstedt) and stored at 4°C prior to use in plaque assays. Plaque assays were undertaken within 24 hr of collecting these samples. Phages were initially isolated and enumerated using a simple single layer plaque assay (67). However, where this was unsuccessful a modified plaque assay was used that allowed a greater volume of water to be added. Briefly, filtered seawater was mixed with CaCl_2_ to a final concentration of 1 mM followed by addition of *E. coli* MG1655 cells at a 1:20 ratio and incubating the mixture at room temperature for 5 minutes. Subsequently, samples were mixed with molten LB agar at a 1:1 ratio, final agar concentration 0.5% (w/v). Agar plates were incubated overnight at 37°C and checked for the presence of plaques. For samples in which no coliphage were detected an enrichment procedure was carried out. Briefly, 20 mL filtered seawater was mixed with 20 mL LB broth and 1 mL *E*. *coli* MG1655 (OD600=~0.3 i.e. mid-exponential phase) and incubated overnight at 37°C, followed by filtration through a 0.22 µm pore-size filter. Phages from this enriched sample were then isolated using the standard plaque assay procedure. Three rounds of plaque purification were used to obtain clonal phage isolates (67).

### Genome Sequencing

Phage DNA was prepared using a previously established method (68). DNA was quantified using Qubit and 1 ng DNA used as input for NexteraXT library preparation following the manufacturer’s instructions. Sequencing was carried out using a MiSeq platform with V2 (2 x250 bp) chemistry. Fastq files were trimmed with Sickle v1, using default parameters (69). Genome assembly used SPAdes v3.7 with the careful option (70). Reads were then mapped back against the resulting contig with BWA MEM v0.7.12 (71) and SAM and BAM files manipulated with SAMtools v1.6 to determine the average coverage of each contig (71). If the coverage exceeded 100x then the reads were subsampled and the assembly process repeated, as high coverage is known to impede assembly (68). Phage genomes were then annotated with Prokka using a custom database of all phage genomes that had previously been extracted from Genbank (72). Further annotation was carried out using the pVOG database to annotate any proteins that fall within current pVOGS using hmmscan (73, 74). Raw sequence data and assembled genomes were deposited in the ENA under the project accession number PRJEB28824

### Bioinformatics and comparative genomics

A MASH database was constructed of all complete bacteriophage genomes available at the time of analysis (~ 8500, April 2018) using the following mash v2 settings “ –s 1000” (75). This database was then used to identify related genomes based on MASH distance which has previously been shown to be equivalent to ANI (75). Phage genomes that were found to be similar were re-annotated with Prokka to ensure consistent gene calling between genomes for comparative analysis (72). Core genome analysis was carried out with ROARY using “--e --mafft -p 32 –i 90” as a starting point for analysis (76). These parameters were adjusted as detailed in the text. The optimal phylogenetic markers were determined using the GET_PHYLOMARKERS pipeline, with the following settings “-R1 – t DNA” (47). Average nucleotide identity was calculated using autoANI.pl (77). Phylogenetic analysis was carried out using IQ-TREE (78), with models of evolution selected using modeltest (79); trees were visualised in ITOL (80).

### One-step growth experiments

Phage growth parameters (burst size, eclipse and latent period) were determined by performing one-step growth experiments as described by Hyman and Abedon (81), with free phages being removed from the culture by pelleting the host cells via centrifugation at 10,000 g for 1 min, removing the supernatant and resuspending cells in fresh medium (81). Three independent replicates were carried out for each experiment.

### TEM

Representative phages, as determined from genome sequencing, were imaged using a Transmission electron microscope (TEM) as follows: 10 µl high titre phage stock was added to a glow discharged formvar copper grid (200 mesh), left for 2 mins, wicked off and 10 µl water added to wash the grid prior to being wicked off with filter paper. 10 µl 2% (w/v) uranyl acetate stain was added to the grid and left for 30 secs, prior to its removal. The grid was air dried before imaging using a JEOL JEM-1400 TEM with an accelerating voltage of 100kV. Digital images were collected with a Megaview III digital camera using iTEM software. Phage images were processed in ImageJ using the measure tool and the scale bar present on each image to obtain phage particle size (82). Measurements are the average of at least 10 phage particles.

### Preparation of viral proteomes for nanoLC-MS/MS and data analysis

Prior to proteomics high titre phage stocks were purified using CsCl density gradient centrifugation at 35,000 g for 2 hrs at 4 °C. Subsequently, 30 µl concentrated phage was added to 10 µl NuPAGE LDS 4X sample buffer (Invitrogen) heated for 5 min at 95°C and analysed by SDS-PAGE as described (83). Polyacrylamide gel bands containing all phage proteins were excised and standard in-gel reduction with iodoacetamide and trypsin (Roche) proteolysis was performed prior to tryptic peptide extraction (83). Samples were separated and analysed by means of a nanoLC-ESI-MS/MS using an Ultimate 3000 LC system (Dionex-LC Packings) coupled to an Orbitrap Fusion mass spectrometer (Thermo Scientific) with a 60 minute LC separation on a 25 cm column and settings as described previously (83). Compiled MS/MS spectra were processed using the MaxQuant software package (version 1.5.5.1) for shotgun proteomics (84). Default parameters were used to identify proteins (unless specified below), searching an in-house-generated database derived from the translation of phage genomes. Firstly, a six reading frame translation of the genome with a minimum coding domain sequence (CDS) cut-off of 30 amino acids (*i.e.* stop-to-stop) was used to search for tryptic peptides. Second, the search space was reduced by using a database containing only CDS detected in the first database search, again, looking for tryptic peptides. Finally, the reduced CDS database was also searched using the N-terminus semi-tryptic digest setting to find the protein N-terminus. Analysis was completed using Perseus software version 1.6.0.7 (85). All detected peptides from all three analyses are compiled in Supplementary Table S5. Only proteins detected with two or more non-redundant peptides were considered.

## Supporting information

## Acknowledgements

Bioinformatic analysis was carried out using MRC CLIMB Infrastructure MR/L015080/1. AM was funded by NERC AMR-EVAL FARMS (NE/N019881/1). T.R. and S.M. were in receipt of PhD studentships funded by the Natural Environment Research Council (NERC) CENTA DTP. A.G. was in receipt of a PhD studentship funded by the Engineering and Physical Sciences Research Council (ESPRC) SynBio

---

**Table S1.** Core-genes, ANI and genes used for phylogenetic analysis of phages within the genus *RB69virus.* All phages were re-annotated to ensure consistent gene calling. ANI was calculated using autoANI.

**Table S2.** Core-genes, ANI, and genes used for phylogenetic analysis of phages within the genus *T5*virus. All phages were re-annotated to ensure consistent gene calling. ANI was calculated using autoANI.

**Table S3.** Core-genes, ANI, and genes used for phylogenetic analysis of phages within the subfamily *Tunavirinae.* ANI was calculated using autoANI.

**Table S4.** Host range of coliphages vB_Eco_swan01, vB_Eco_mar005P1, vB_Eco_mar002J2, vB_Eco_mar003J3 and vB_Eco_mar004NP2 against Enterobacteriaceae hosts. Infected hosts are marked with a black box and those that are not infected with

| Host bacterial strains | Phage Isolate |  |  |  |  |
| --- | --- | --- | --- | --- | --- |
|  | vB_Eco_mar002J2 | vB_Eco_mar003J3 | vB_Eco_mar004NP2 | vB_Eco_mar005P1 | vB_Eco_swan01 |
| *Escherichia coli* MG1655 (K12) | 1 | 1 | 1 |  | 1 |
| *Escherichia coli* GD45 | - | - | - | - | - |
| *Escherichia coli* GU48 | - | - | 1 | - | - |
| *Escherichia coli* T3-21 | - | - | - | - | - |
| *Escherichia coli* SFR-11 | - | - | - | - | - |
| *Escherichia coli* D22 | 1 | 1 | 1 | - | 1 |
| *Escherichia coli* N43 | 1 | 1 | 1 | 1 | 1 |
| *Escherichia coli* EV36 | 1 | 1 | 1 | 1 | 1 |
| *Escherichia coli* 170713 | - | - | - | - | - |
| *Escherichia coli* 170972 | - | - | - | - | - |
| *Klebsiella varicola* DSM 15968 | - | - | - | - | 1 |
| *Klebsiella oxytoca* DSM 5175 | - | - | - | - | - |
| *Klebsiella oxytoca* DSM 25736 | - | - | - | - | - |
| *Klebsiella quasipneumoniae* DSM28211 | - | - | - | - | - |
| *Klebsiella michiganensis* DSM 25444 | - | - | - | - | - |
| *Klebsiella pneumoniae pneumoniae* DSM30104 | - | - | - | - | - |
| *Klebsiella pneumoniae* isolate 170723 | - | - | - | - | - |
| *Klebsiella oxytoca* isolate 170748 | 1 | - | 1 | - | - |
| *Klebsiella pneumoniae* isolate 170820 | - | 1 | - | - | - |
| *Klebsiella oxytoca* isolate isolate 170821 | - | - | - | - | - |
| *Klebsiella pneumoniae* isolate 170958 | 1 | 1 | 1 | - | - |
| *Klebsiella pneumoniae* isolate 171167 | 1 | 1 | - | - | - |
| *Klebsiella oxytoca* isolate 171266 | - | - | - | - | - |
| *Klebsiella pneumoniae* isolate 170304 | - | - | - | - | - |
| *Salmonella typhimurium* | - | - | 1 | - | - |

**Table S5** Peptides detected from phages vB_Eco_swan01, vB_Eco_mar005P1, vB_Eco_mar002J2, vB_Eco_mar003J3 and vB_Eco_mar004NP2.

## Supplementary Figures

**Figure S1.** Phylogenetic analysis of phages within the genus *RB69virus.* The tree is based on the nucleotide sequence of the terminase gene, using a TIM2+F+R5 model of evolution, with 1000 bootstrap replicates using IQTREE (78).

**Figure S2.** Phylogenetic analysis of phages within the genus *T5virus.* The phylogenetic tree is based on the nucleotide sequence of the terminase gene, using a TIM2+F+R3 model of evolution, with 1000 bootstrap replicates using IQTREE (78).

**Figure S3.** Phylogenetic analysis of phages within the subfamily *Tunanvirnae.* The tree is based on the nucleotide sequence of the terminase gene, using a TIM2+F+R3 model of evolution, with 1000 bootstrap replicates using IQTREE (78).

**Figure S5 A)** The abundance of representative bacteriophages in the Baltic Virome (DNA). Genome coverage was calculated using BBMap with the following options ‘covstats minid=90` using the Baltic Virome fasta data available from iMicrobe under project code CAM_P_0001109. The coverage data presented was calculated by the covstats function within BBMap. **B)** Abundance of transcripts from representative bacteriophages from the Baltic metatranscriptomic dataset. Reads from the metatranscriptomics dataset were sequentially downloaded using fasterq-dump from the short read archive. Reads were again stringently mapped to a single file that contained all representative genomes using BBMap with the settings ‘minid=90, covstats, outm’. The number of reads mapped to each genome was normalised for both length of the phage genome and the number of reads per sample.

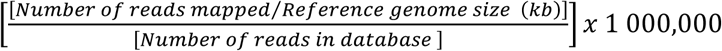

To give the number of reads mapped per kb of phage genome per million reads in the database. This data was plotted for each sampling site. For ease of display only accession numbers are plotted as used in the original publication for this data (56).

