## Supplementary figures and images for "Riding the wave of genomics, to investigate aquatic coliphage diversity and activity"

### Supplementary file 1

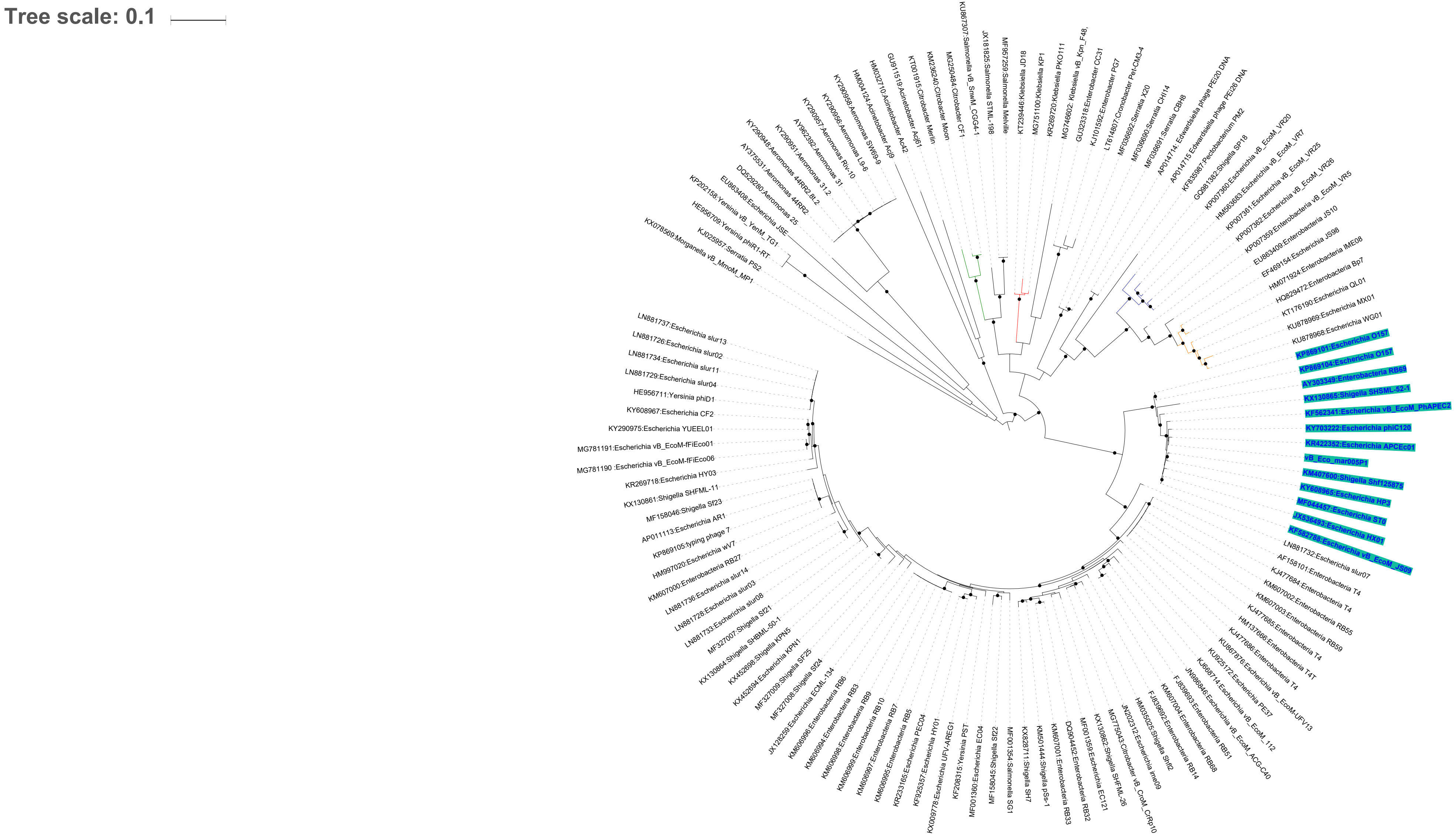

### Supplementary file 2

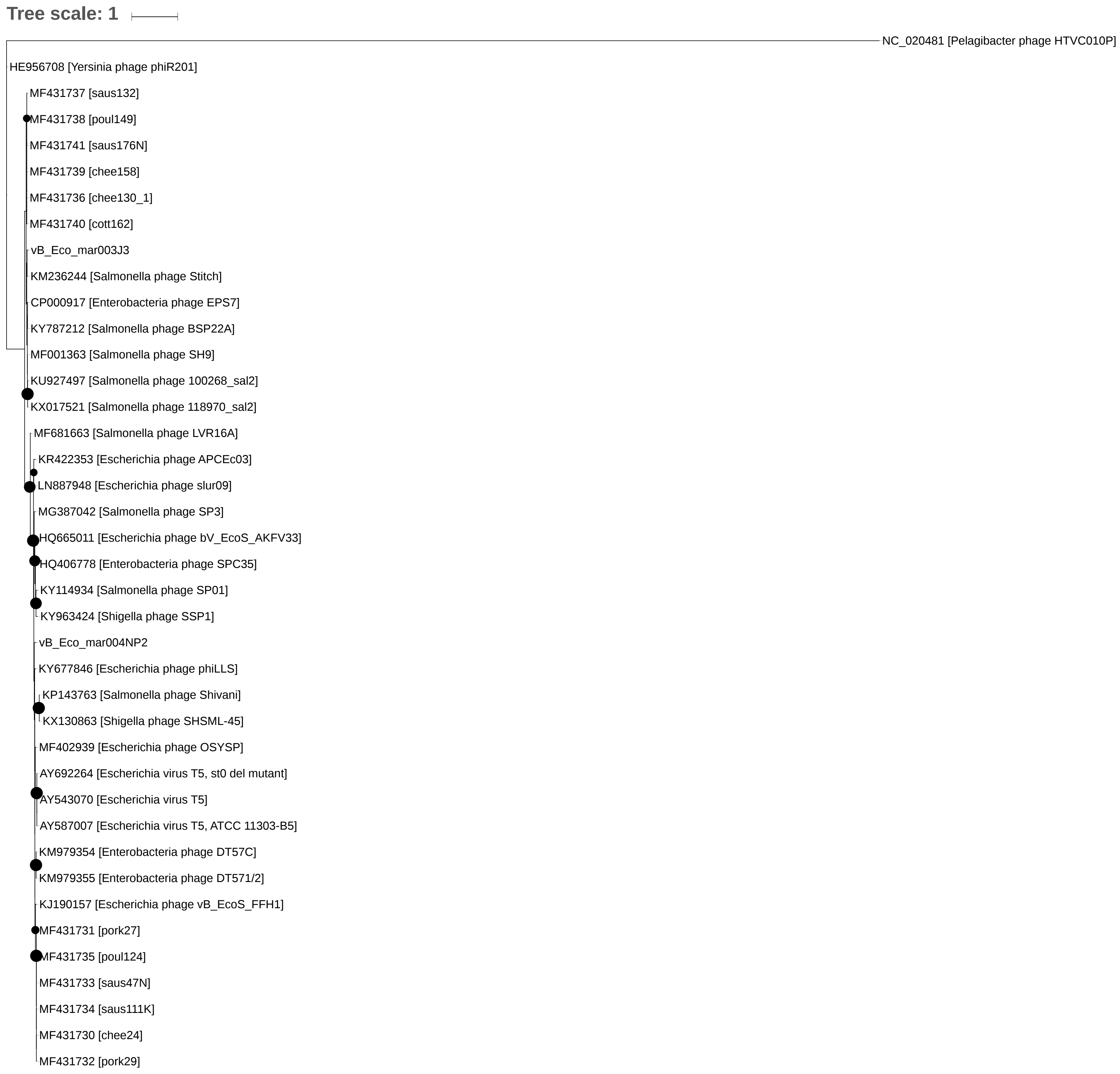

### Supplementary file 3

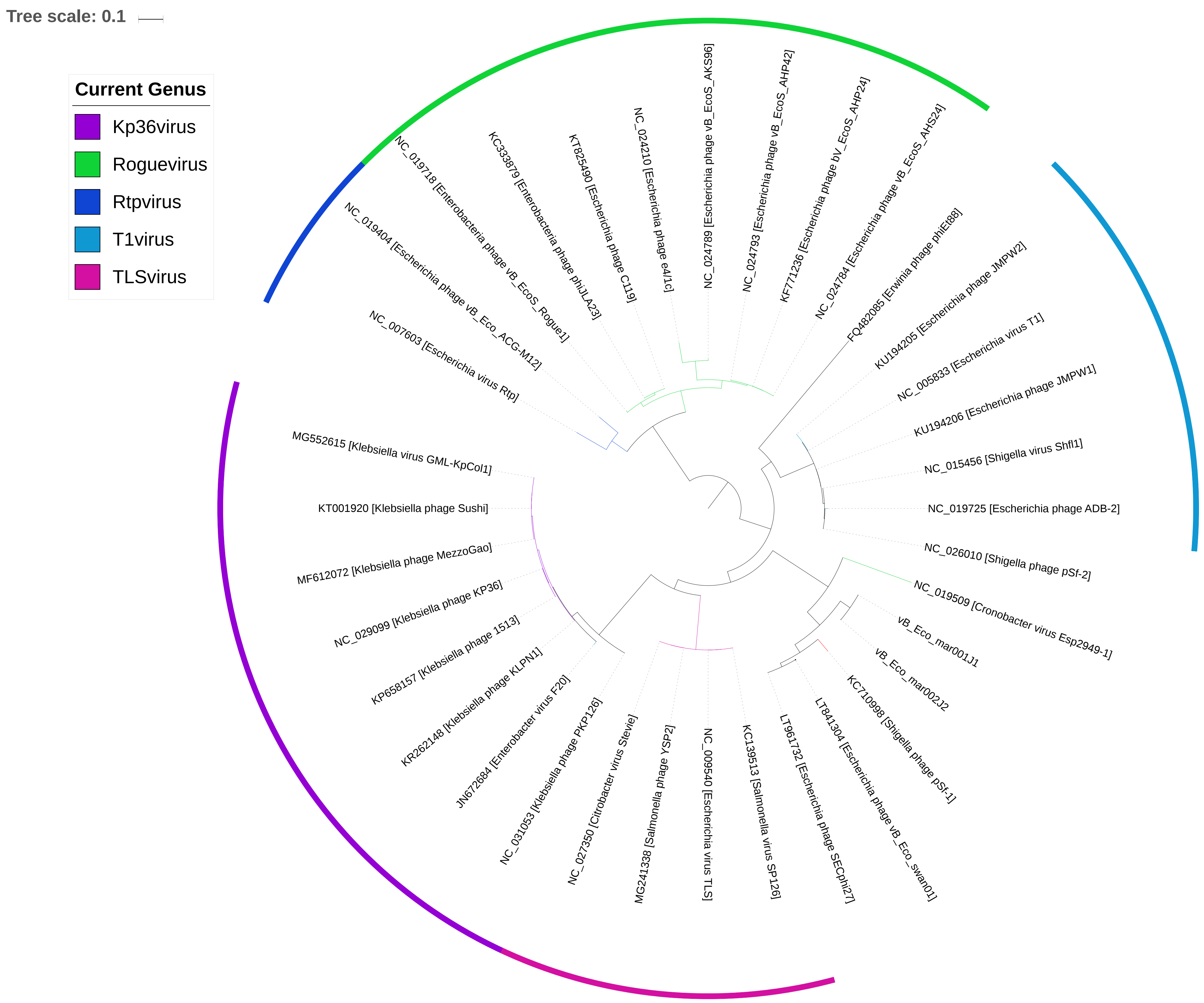

### Supplementary file 5

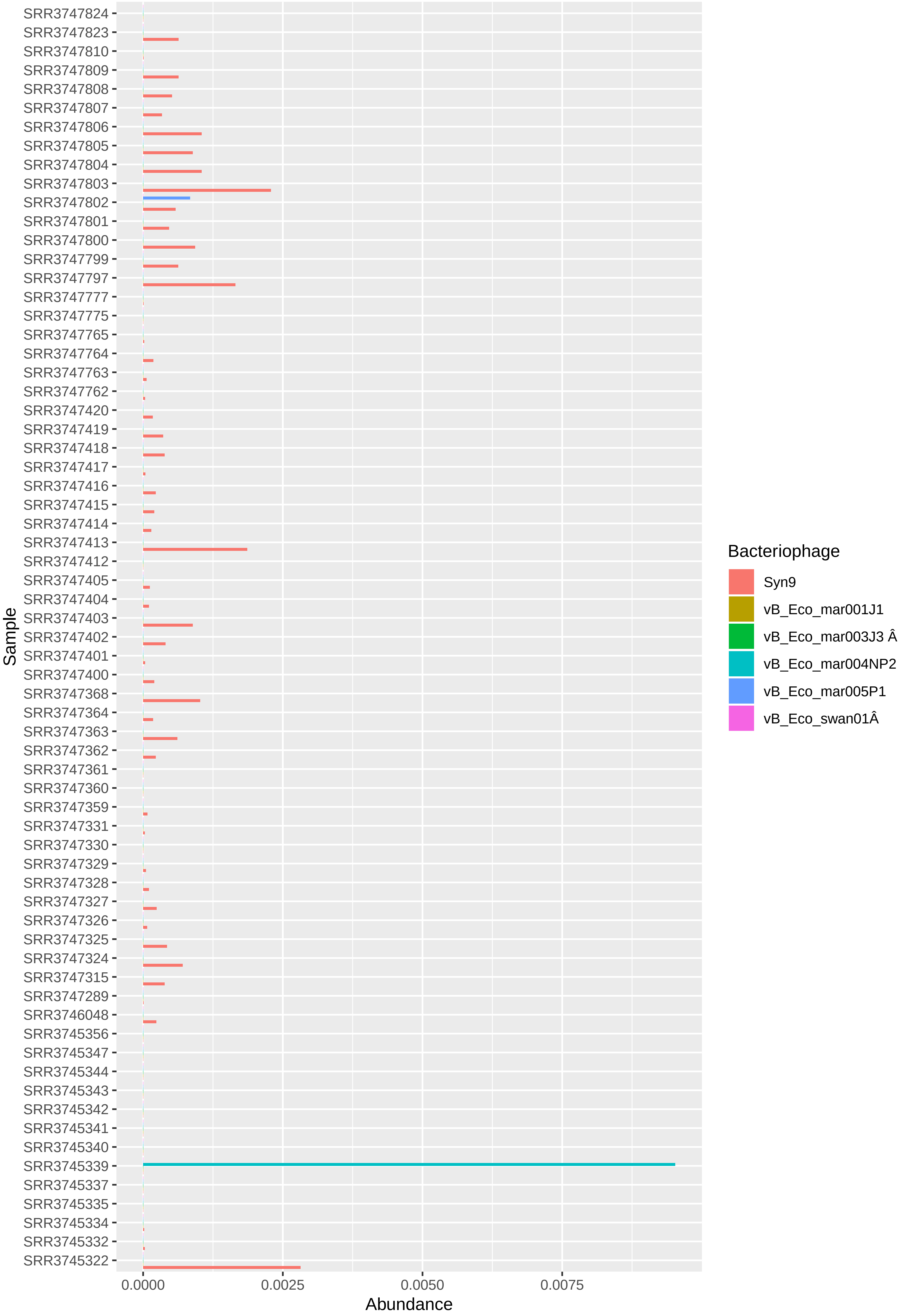
