## Supplementary material for "Riding the wave of genomics, to investigate aquatic coliphage diversity and activity"

### Slide 1
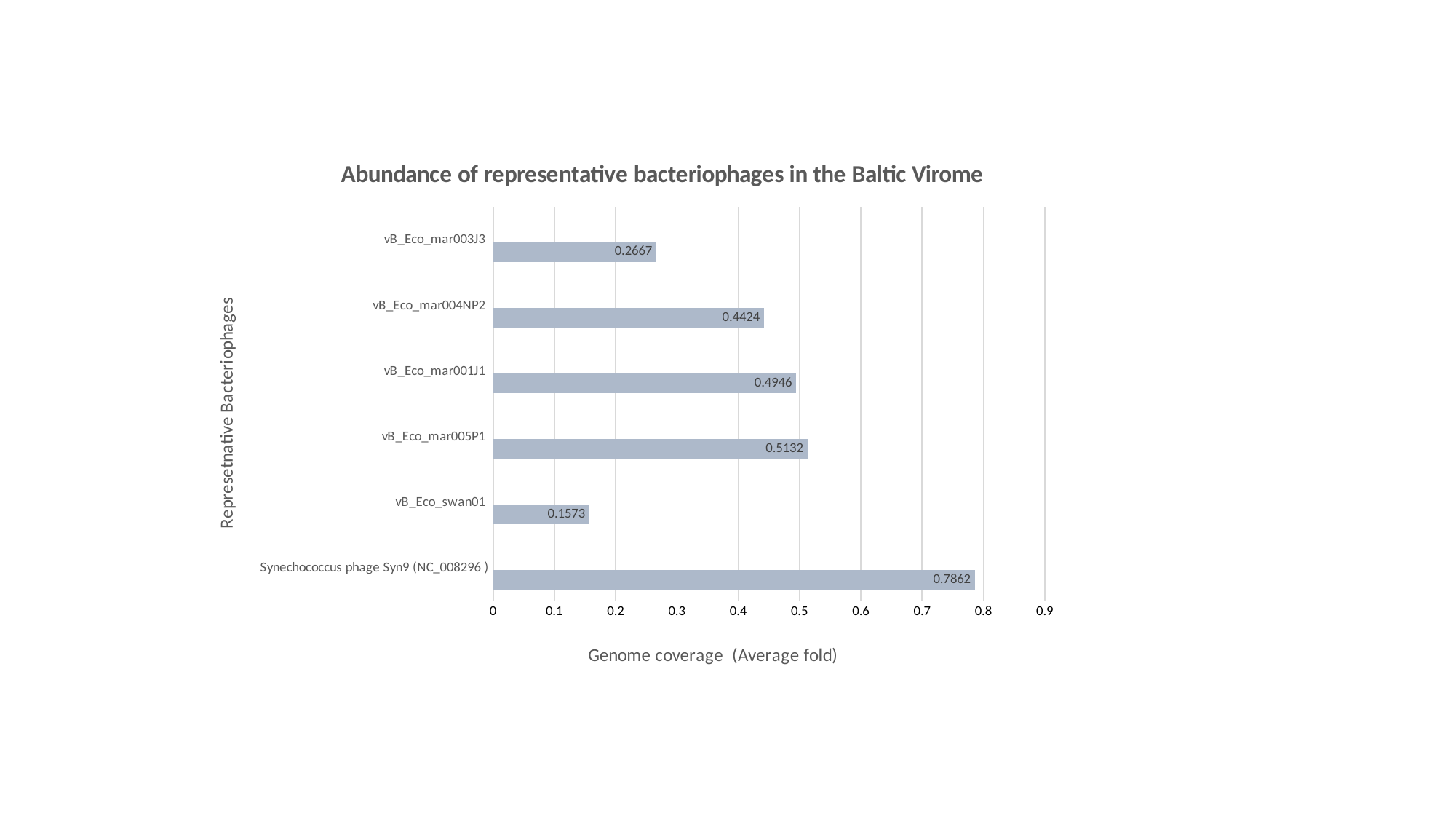

#### Chart: Abundance of representative bacteriophages in the Baltic Virome
| Category | Covered_percent | |
|---|---|---|
| Synechococcus phage Syn9 (NC_008296 ) | 0.7862 | None |
| vB_Eco_swan01  | 0.1573 | None |
| vB_Eco_mar005P1 | 0.5132 | None |
| vB_Eco_mar001J1 | 0.4946 | None |
| vB_Eco_mar004NP2 | 0.4424 | None |
| vB_Eco_mar003J3 | 0.2667 | None |
